# Collaborations between CpG sites in DNA methylation

**DOI:** 10.1101/149815

**Authors:** You Song, Honglei Ren, Jinzhi Lei

## Abstract

DNA methylation patterns have profound impacts on genome stability, gene expression, and development. The molecular base of DNA methylation patterns has long been focused at single CpG sites level. Here, we construct a kinetic model of DNA methylation with collaborations between CpG sites, from which a correlation function was established based on experimental data. The function consists of three parts that suggest three possible sources of the correlation: movement of enzymes along DNA, collaboration between DNA methylation and nucleosome modification, and global enzyme concentrations within a cell. Moreover, the collaboration strength between DNA methylation and nucleosome modification is universal for mouse early embryo cells. The obtained correlation function provide insightful understanding for the mechanisms of inheritance of DNA methylation patterns.

## 1. Introduction

DNA methylation is a process by which methyl groups are added to DNA segments. Methylation represents a key epigenetic modification that changes the activity of a DNA segment without changing the sequence. The DNA methylation patterns are dynamically regulated during development, and are important for stable silencing of gene expression, maintenance of genome stability, and establishment of genomic imprinting ^1,18^. Methylation of the fifth position of cytosine (5-methylcytosine, 5mC) is found in most plant, animal, and fungal models, and is primarily restricted to palindromic CpG (CG/GC) dinucleotides ^12,16^.Biologically, the molecular base of DNA methylation patterns has been focused at *single* CpG sites level, which is regulated by various types of enzymes, such as DNMT1/UHRF1 for the maintenance of methylation marks, TET families for the active demethylation, and DNMT3a/DNMT3b for the *de novo* methylation ^18^.

There are now substantial evidences that additional mechanisms should be at work to ensure the robust maintenance of methylation patterns. During mouse gametes and early embryos, most functional genomic elements undergo significant demethylation, except CpG islands (CGIs) and 5’ untranslated regions (UTRs) whose methylation levels are already very low in gametes ^17^. In zebrafish early embryos, the oocyte methylome is gradually discarded after 16-cell stage, and then progressively reprogrammed to a pattern similar to that of the sperm methylome 10. DNA methylation patterns of human hematopoietic stem cells (HSCs) from four different sources show different profiles, and DNA methylation dynamics of myeloid-lymphoid lineage choice display an asymmetric pattern ^6^. These collaborative changes in DNA methylation patterns suggest potential correlations between the kinetics of CpG sites methylation/demethylation.

A computation model has shown that dynamic collaboration between nearby CpG sites can provide strong error-tolerant inheritance of methylation states of a cluster of CpGs, and shown stable bimodal methylation patterns ^8^. It was proposed that the collaboration can be achieved by recruitment of methylase or demethylase enzymes. However, the molecular based of the correlation remains unclear, and this proposed mechanism fail to explain the CpG site distance dependence correlation from experimental data (Figure 1b).

**Fig. 1.**
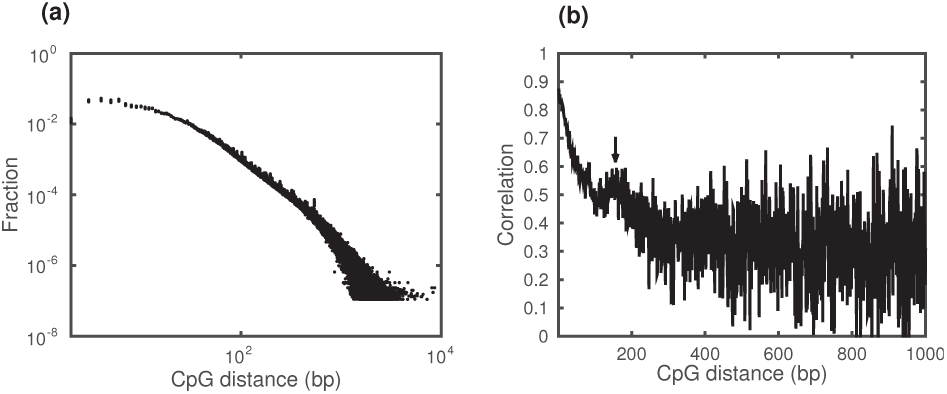
Experimental data. (a) Distribution of the distances between adjacent CpG sites from chr1 to chr19 of mouse DNA. (b) Pearson correlations between DNA methylation levels of adjacent CpG sites of given distance (in bp). Data obtained from the parental strain chromosome 1 of 2-cell stage at mouse embryo (P-chr1-2-cell-mouse) (data source: GSM1386021).

In this study, we examine experimental data for patterns of DNA methylation correlation between adjacent CpG sites, and establish a computational model of methylation/demethylation kinetics in which CpG site distance dependence correlations are involved. The correlation function is obtained from experimental data. Mechanisms of correlation are discussed in accordance with the obtained correlation function.

## 2. Results

### 2.1. Pearson correlations between CpG site methylations

To examine the patterns of DNA methylation, we study data from mouse early embryo (GSM1386021) ^17^, which includes 5mC reads of each CpG sites from different stages of mouse embryo from sperm/oocyte to E13.5. In mouse genome, the distances between adjacent CpG sites range from 2 to a few thousand bp. Different chromosomes show similar spectrum of CpG distance distributions and the frequency decreases with the CpG distance (Figure 1a). For adjacent pairs of CpG sites with given distance *d* (in bp), the Pearson correlation *C*(*d*) was obtained as
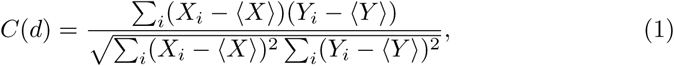

where (*X_i_, Y_i_*) is the DNA methylation level of a pair of CpG sites, 〈*X*〉 and 〈*Y*〉 are average of {*X_i_*} and {*Y_i_*}, respectively.

We examine data from the parental strain chromosome 1 of 2-cell stage at mouse embryo (P-chr1-2-cell-mouse in short), and see clear correlation between methylation levels of adjacent CpG sites. There is strong correlation with Pearson’s *r* = 0.84 when the CpG distance is 2 bp, and the correlation decreases with the CpG distance (Figure 1b). The correlation displays a local maximum at the distance of about 160 bp (Figure 1, arrow). Moreover, the Pearson correlation is consistent when we examine data for different chromosomes in the same cell (Appendix A). While we calculate the correlation patterns from cells of different embryonic stages, the correlations show evolutionary dynamics during embryo development (Appendix B). These results suggest that the correlation pattern is universal for all chromosomes, however can be dependent on the cellular status.

### 2.2. Kinetic model of DNA methylation

To investigate the molecular mechanism of correlation between CpG sites, we construct a kinetic model aiming to uncover the molecular mechanism of CpG sites correlation in DNA methylation. Here, we introduce a key assumption that the correlation between methylation/demethylation of CpG sites is dependent on the CpG distance (Figure 2). Each CpG site can transit between fully un-metheylated (u), half-methyalted (h), and fully methylated (m) states (solid arrows in Figure 2). The transition rates are affected by the status of nearby CpGs (dashed arrows in Figure 2), and the correlation strength *ϕ*(*d*) is represented with a function of the distance (in bp) between the two CpGs. Our aim was to formulate the function *ϕ*(*d*) to reproduce the experimental correlation.

**Fig. 2.**
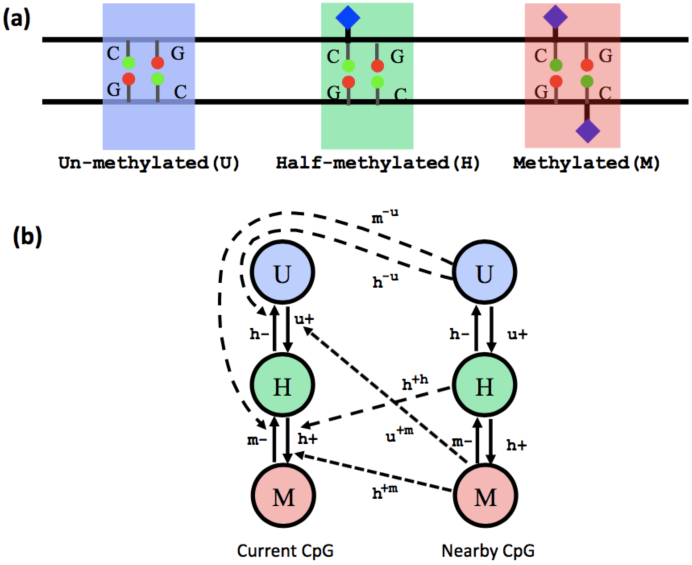
Schematic illustration of the model. (a) Three possible states of a CpG: un-methylated (U), half-methylated (H), or methylated (M). (b) Kinetics of the transitions between the methylation state of a CpG site. Solid arrows represent the transition between the states, dashed arrows show the correlations from the nearby CpG.

Our model was established based on the collaboration model in Haerter et al. (2014)^8^. The *i*’th CpG site can at one of the states u, h, or m, and randomly methylate with a rate 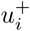 from the state u to h, or a rate 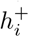 from the state h to m. Moreover, the methyl mark can be removed from the CpG site with a rate 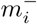 from m to h, or a rate 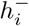 from h to u. The kinetic rates 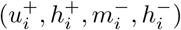 consist of two parts of contributions, the contribution from basal level rates (*u*^+^, *h*^+^, *m*^−^, *h*^−^), and the contribution due to collaboration between neighboring CpG sites. According the assumptions in Haerter et al. (2014)^8^, when a CpG is at state m, it promotes the methylation of the nearby CpG, and hence the methylation rate from u to h increases by *ϕ*(*d*)(*u^+m^*−*u^+^*), and the rate from h to m increases by *ϕ*(*d*)(*h^+m^*−*h^+^*). Similarly, a CpG of state h contributes an increase *ϕ*(*d*)(*h^+h^*−*h^+^*) to the methylation rate of the nearby CpG site 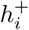, and a CpG of state u promotes the demethylation processes, so that the demethylation rate 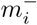 increases by *ϕ*(*d*)(*m*^−*u*^ − *m*^−^), and the rate 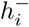 increases by *ϕ*(*d*)(*h*^−*u*^ − *h*^−^).

Mathematically, the kinetic rates of the *i*’th CpG in a DNA sequence are given by
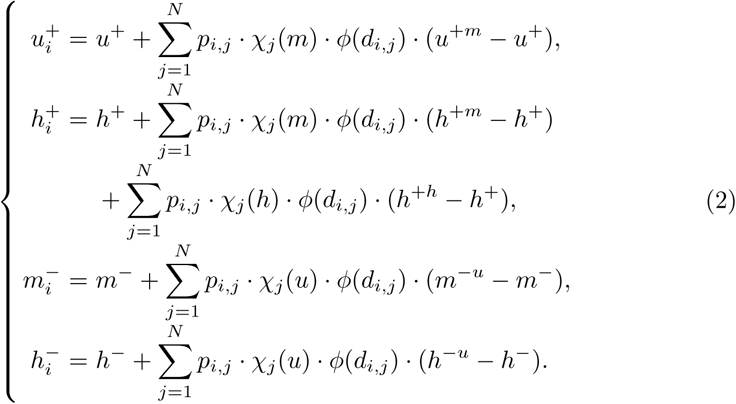

Here *p_i,j_* = 1 when the *i*’th CpG is affected by the *j*’th CpG, otherwise *p_i,j_* = 0. The index *χ_j_* (*s*) = 1 (*s* = *u, h, m*) if the *j*’th CpG has the status *s*, and *χ_j_* (*s*) = 0 if otherwise. We simply consider the nearby correlation so that *p_i,j_* = 1 only when *j* = *i* ± 1. Here we note that although only the nearby CpGs are involved in the correlation, however the distance between nearby CpG sites can be separated by hundreds base pairs. We note that the function *ϕ*(*d*) is assumed to be the same for all type correlations. This simplification was trying to construct a minimal model, however biologically can be different. In model simulations, we refer the kinetic models from Haerter et al. (2014)^8^, which are listed at Table 1.

The kinetic rates in Eq. 2 define the dynamics of in each cell cycle. To include the effect of reallocation at DNA replication, we reset the status *m* to *h* at the beginning of each cell generation. For a given DNA sequence and a well defined function *ϕ*(*d*), stochastic simulation based on the above kinetic rates for ~20 generations lead to a stationary DNA methylation pattern, which give a model predicted correlation profile.

**Table 1.** Default kinetic parameter values used in the simulations. Parameters are referred to Haerter et al.(2014).

| $u^+$ | $h^+$ | $m^-$ | $h^-$ | $h^{+h}$ | $h^{+m}$ | $u^{+m}$ | $h^{-u}$ | $m^{-u}$ |
| --- | --- | --- | --- | --- | --- | --- | --- | --- |
| 0.008 | 0.008 | 0.04 | 0.04 | 0.24 | 0.24 | 0.24 | 0.05 | 0.05 |

### 2.3. Correlation function

To obtain the function *ϕ*(*d*), we refer the experimental data from P-chr1-2-cell-mouse, denoted as C(d) (Figure 1b). The procedure is given below.

1. Set *ϕ* in model as a constant independent to the distance *d* and run the model simulation. Thus, the correlation defined in above is dependent on the value of *ϕ*, however is independent to the distance *d* (Figure 3(a)). Larger *ϕ* leads larger correlations (Figure 3(b)). Fitting the dependence of the simulated correlation with *ϕ*, we obtained the function *R*_0_(*ϕ*) (Figure 3(b), red)
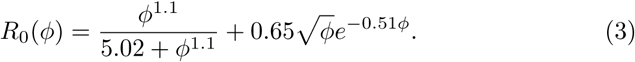 In fitting the data, we first fit with Hill type function 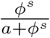 to obtain good fit with *ϕ <* 5 to have *s* = 1.1, next fit the remaining part of data with the function of form 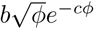 to obtain *c* = 0.51. Finally, we adjust the coefficients *a* and *b* to obtain a good fit for overall data. The function *R*_0_(*ϕ*) is monotonous increasing.
2. Consider the simulated correlation *R*_0_(*ϕ*) and the correlation obtained from experimental data (Figure 1), which is denoted as *C*(*d*). While assuming the two correlations equal to each other, *i.e.*, *R*_0_(*ϕ*) = *C*(*d*), we obtain the approximation 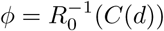 (Figure 3(c)).
3. Substitute the function *ϕ*(*d*) obtained at the above step and run the model simulation to obtain the first approximation of correlation, which is denoted as *C*_0_(*d*) (Figure 3(d)). We note that the simulated correlation *C*_0_(*d*) has the same tendency as the experimental data *C*(*d*), however *C*_0_ (*d*) is in general larger than *C*(*d*).
4. Let the correlation *R*_0_(*ϕ*) equals *C*_0_(*d*), which gives *R*_0_(*ϕ*) = *C*_0_(*d*). Solve this equations to have the correlation function 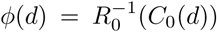 (Figure 3(e), blue dots). We note that the numerical data show power law decay in small *d* region, and a bell-shape bulb at around *d* = 160. Hence, we introduce a function that combines power law decay 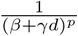, a bell-shape bulb function 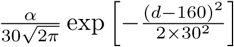, and a constant tail, so that the the numerical data can be fitted with a correlation function of form (Figure 3(e), red line):
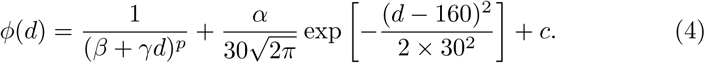 In numerical fitting, we can first find the power law *p* with data of small *d* region (*d <* 130), and then find the fitting coefficients through `FindFit` of `Mathematica`.
5. Now, we substitute the correlation function (4) into model equations, and run the simulation to obtain the simulated correlation, which show good agreement with experimental results (Figure 3(f)).

**Fig. 3.**
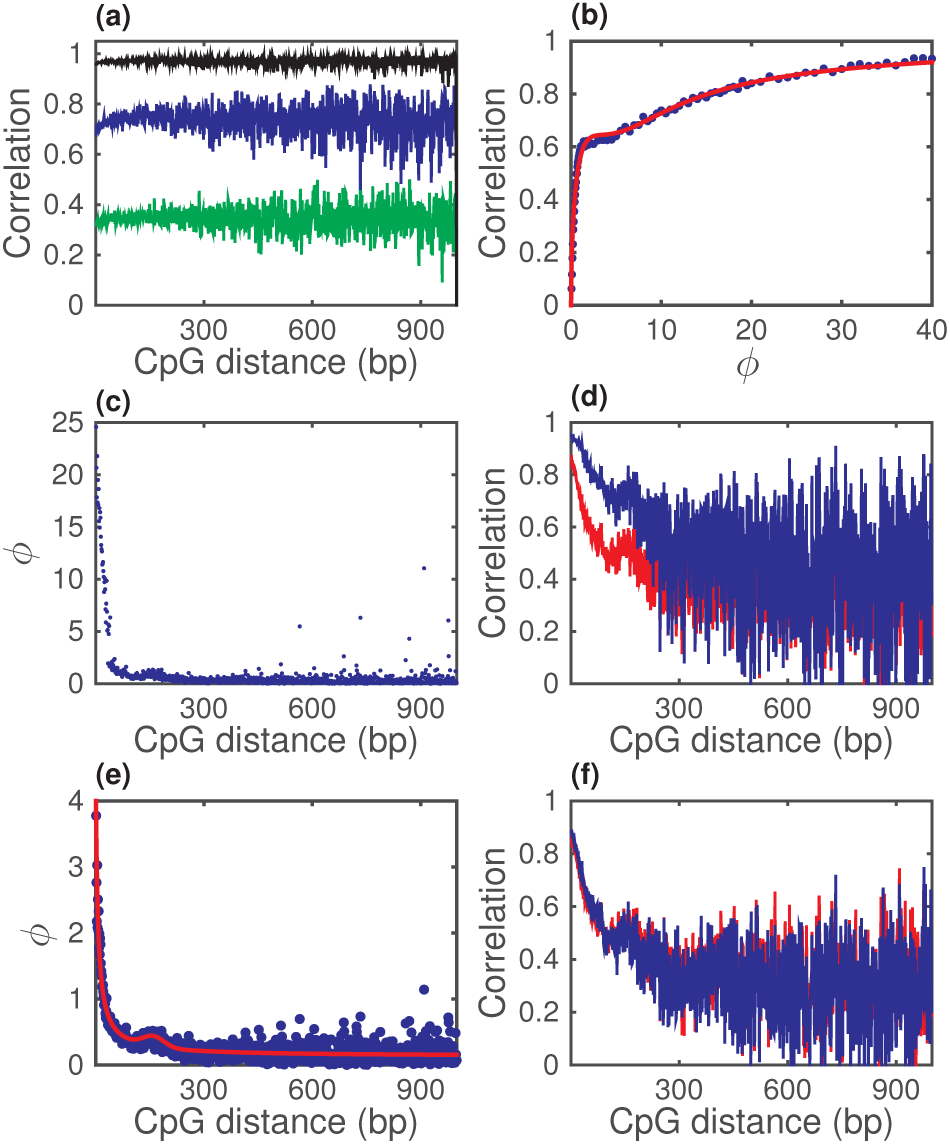
Procedure to obtain the correlation function *ϕ*(*d*). (a) Simulated Pearson correlations with fixed value *ϕ: ϕ* = 0.3 (green), *ϕ* = 2 (blue), and *ϕ* = 30 (black). (b) Dependence of the Pearson correlation on the value *ϕ.* Red curve shows the fitting with equation (3). (c) The approximation correlation function calculated by 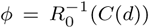. (d) The Pearson correlation obtained from experimental data (red) and from model simulation with 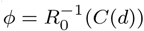 (blue). (e) The correlation function 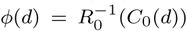 (blue dots). Red curve shows the fitting with equation (4), with parameters *α* = 12, *p* = 1, *β* = 0.195, *γ* = 0.036, *c* = 0.13. (f) Pearson correlations obtained from experimental data (red) and from model simulation with *ϕ*(d) given by equation (4) (blue).

The above procedure gives the correlation function obtained from experimental data. Substituting the obtained *ϕ*(*d*) into Eq. 2 and perform stochastic simulation, we can nicely reproduce the correlation profile, in good agreement with experimental data (Fig. 3f).

We note that both kinetic parameters in Table 1 and experimental data are involved in the procedure of obtaining *ϕ*(*d*). The kinetic parameters usually affect the mean methylation level, and may alter the coefficients in the correlation function (detailed below). However, the formulation of the function *ϕ*(*d*) maintains for different sets of parameters. Here, as we intended to study the correlation, we always fixed the kinetic parameters and examined the dependence of *ϕ*(*d*) on experimental data. Nevertheless, the kinetic parameters are dependent on the enzyme activities, and can be alter from cell to cell.

We apply the procedure to experimental data from different stages of mouse early embryo. The function Eq. (4) is well fitting with different stages data, however the coefficients can be varied (Table 2)(Figure 5). Moreover, we apply the procedure to data from different tissues of mouse and human, and show small root-mean-square deviations (RMSDs) between experimental correlation and model simulation correlation (Appendix C, Table C1). These results imply that the correlation *ϕ*(*d*) of form Eq. (4) can provide insights to the dynamics of DNA methylation.

**Table 2.** Parameters for the correlation functions corresponding to different stages and strands from mouse embryo cells.

| Para. | sperm | 2-cell | 4-cell | E6.5 | E7.5 | PGC E13.5 |
| --- | --- | --- | --- | --- | --- | --- |
| $p$ | 1.5 | 1 | 1 | 1 | 1 | 1 |
| $\alpha$ | 12 | 12 | 12 | 12 | 12 | 12 |
| $\beta$ | 0.001 | 0.195 | 0.3 | 0.15 | 0.04 | 0.09 |
| $\gamma$ | 0.0068 | 0.04 | 0.036 | 0.06 | 0.042 | 0.06 |
| $c$ | 0.005 | 0.13 | 0.1 | 0.06 | 0.02 | 0.001 |

| Para. | oocyte | 2-cell | 4-cell | E6.5 | E7.5 | PGC E13.5 |
| --- | --- | --- | --- | --- | --- | --- |
| $p$ | 1.5 | 1 | 1 | 1 | 1 | 1 |
| $\alpha$ | 12 | 12 | 12 | 12 | 12 | 12 |
| $\beta$ | 0.07 | 0.0015 | 0.032 | 0.001 | 0.001 | 0.138 |
| $\gamma$ | 0.0135 | 0.0092 | 0.0065 | 0.05 | 0.03 | 0.075 |
| $c$ | 0.19 | 0.09 | 0.07 | 0.07 | 0.02 | 0.001 |

### 2.4. Effects of kinetic parameters to the correlation function

To examine the effects of changing kinetic parameters to the correlation function, we increase or decrease each kinetic parameter by 50% and examine the corresponding changes in the coefficients in the correlation function. Default values of the kinetic parameters are listed in Table 1, changes in the average methylation level and parameters in the correlation function in response to changes in kinetic parameters, are given by Figure 4. Results show that changing the kinetic parameters can affect the average methylation level, as well as the coefficients *β, γ* and *c* in the correlation function given by Eq. 4 (here we fixed *p* = 1 and *α* = 12).

Usually, decreasing the methyl adding rates *u^+^* and *h^+^* decrease the methylation level, and decreasing the methyl removing rates *m*^−^ and *h*^−^ increase the methylation level. The average methylation level is not sensitive to changes in the correlative parameters *h^+h^, h^+m^, u^+m^, h*^−*u*^, and *m*^−*u*^. These observations are confirmed by Figure 4.

**Fig. 4.**
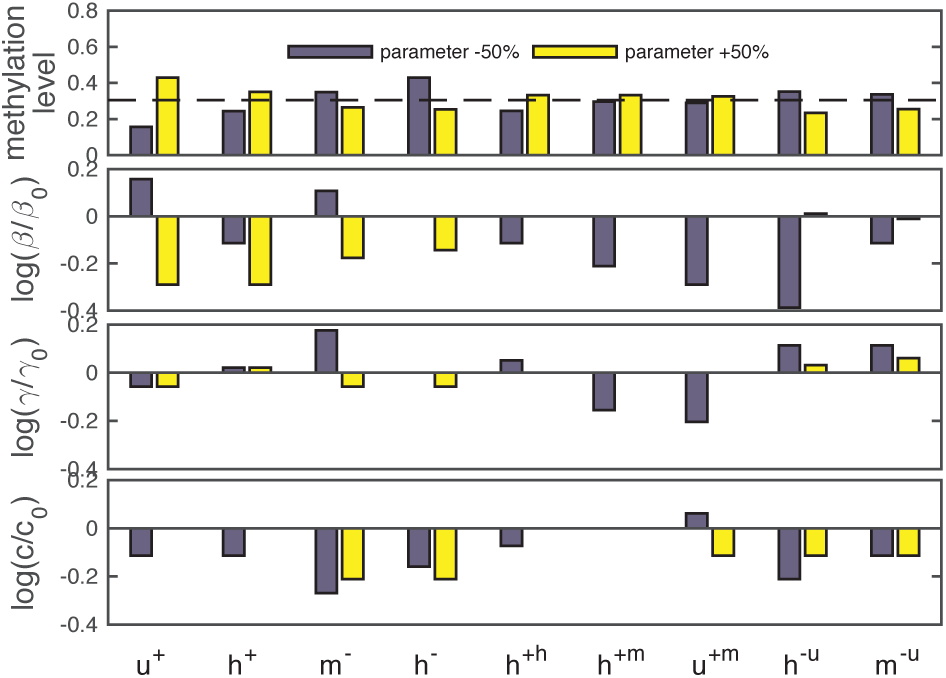
Effects of changes in the kinetics parameters. We either increased (blue bars) or decreased (yellow bars) each kinetic parameters by 50%, and examine the effects to the average methylation level and to the coefficients in the correlation function Eq. 4. Here the dashed line at the upper panel shows the methylation level with default parameters. Here *β, γ, c* represent the coefficients for modified parameters, and *β*_0_, *γ*_0_,*c*_0_ represent the coefficients for default parameters, respectively. We always set *α* = 12 in fitting the data. See the text for discussion. In the last three panels, the absent bars (zero) mean no difference in the parameter values.

From Figure 4, the consequences of changing the kinetic parameters to the correlation function parameters *β, γ*, and *c* are complicated. According to the procedures of obtaining the correlation function, the kinetic parameters mainly affect the function *R*_0_(*ϕ*) and the first order approximation *C*_0_(*d*). However, these two functions are obtained from model simulation, therefore the stochastic effects are non-negligible in obtaining these correlation function parameters.

### 2.5. The parameter ff in the correlation function

For the parameter *α*, we have seen from Table 2 that *α* = 12 can be applied to different stage embryonic cells. This suggests that the correlation may be insensitive with the parameter *α.* To test is hypothesis, we alter the value *α* to examine its effect to the simulated correlations. We increase or decrease *α* by 50% in Eq. 4 and keep other parameters unchanged and run the model simulation. We repeat the simulations based on data of different embryonic cells, and found no obvious variations in obtained correlations (Figure 5). The RMSD of between simulated correlations and correlations obtained from experimental data are not sensitive with the parameter *α* Table 3. We further apply the produce to various mouse and human somatic tissue cells, the results give two catalogues cells, either *α* = 0.001 or *α* = 4 (Appendix C, Table C1). These results suggest that the *α* can be a universal parameter under certain conditions. Nevertheless, the mechanism waiting for further investigation through models with more kinetics details of the regulation of DNA methyaltion.

**Fig. 5.**
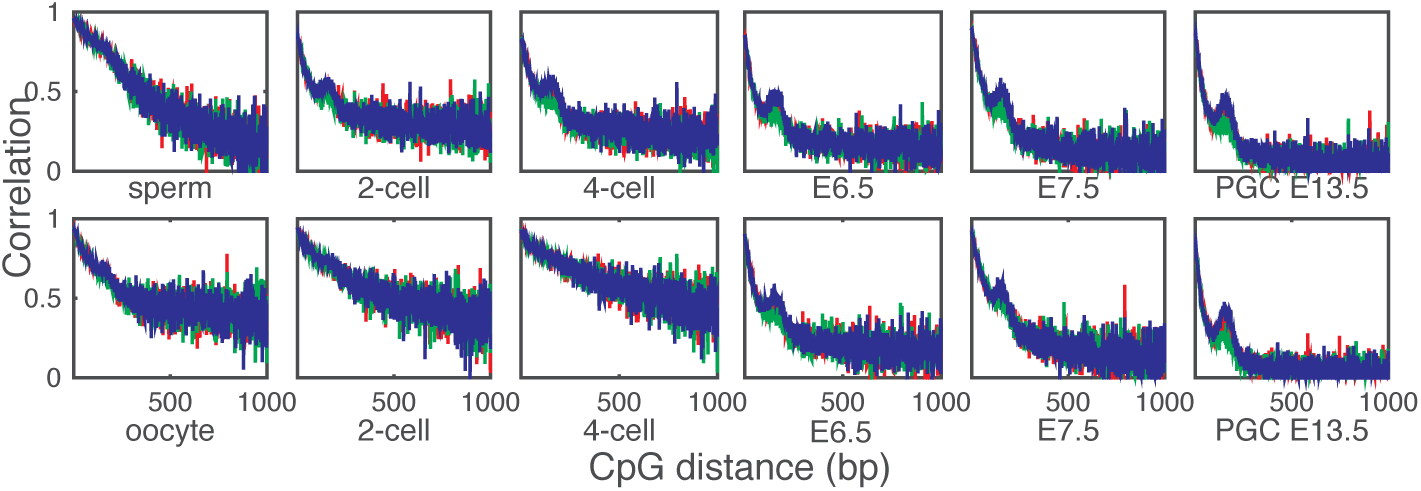
Correlations with *α* = 6.5 (decreased by %50, green), *α* = 12 (default, red) or *α* = 19.5 (increased by %50, blue) based on diaerent stages of mouse embryonic cells. Other coefficients or kinetic parameters were taken as default values as Table 1 and Table 2.

**Table 3.** The Root Mean Squared errors of correlations from simulations (*α* = 6.5,12 or 19.5) relative to the correlations from experimental data of diaerent stages or strains of mouse embryonic cells.

| $\alpha$ | sperm | 2-cell | 4-cell | E6.5 | E7.5 | PGC E13.5 |
| --- | --- | --- | --- | --- | --- | --- |
| 6.5 | 0.209 | 0.13 | 0.156 | 0.203 | 0.214 | 0.269 |
| 12 | 0.211 | 0.129 | 0.147 | 0.197 | 0.215 | 0.26 |
| 19.5 | 0.209 | 0.129 | 0.146 | 0.194 | 0.207 | 0.254 |
| $\alpha$ | oocyte | 2-cell | 4-cell | E6.5 | E7.5 | PGC E13.5 |
| 6.5 | 0.179 | 0.212 | 0.263 | 0.182 | 0.185 | 0.282 |
| 12 | 0.18 | 0.217 | 0.266 | 0.172 | 0.188 | 0.274 |
| 19.5 | 0.183 | 0.218 | 0.263 | 0.17 | 0.184 | 0.271 |

### 2.6. The parameter c in the correlation function

The constant tail represents a global impact of the correlation. We examine the average methylation level versus the constant tail for different stages in mouse embryo cells and tissues. The results show a nonlinear dependence, except the stage of PGC at E13.5 when the DNA methylation is extreme low due to active demethylation (Figure 6). Moreover, we test the data from human tissue cells, which reveal similar form nonlinear dependences (Figure 6). These results suggest that the constant tail in the correlation function is related to the global methylation level, however the molecular mechanism is waiting for further investigation.

**Fig. 6.**
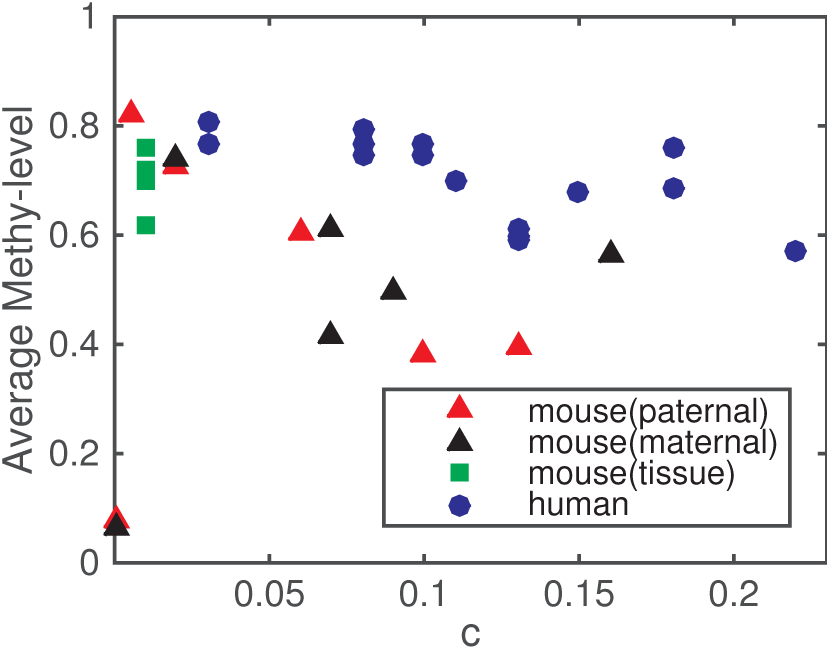
Experimental average methylation level versus constant tail *c* in the correlation function, data obtained form mouse early embryo cells (paternal, red triangles; maternal, blue triangles) and tissue (green squares), or human tissues (blue circles).

## 3. Discussion

We have established a kinetic model of DNA methylation, from which and combining with experimental data we obtained the correlation function for the collaboration effects of adjacent CpG sites methylation. The function O(d) consists of three parts: the power law decay, a bell-shape bulb, and a constant tail. Here we discuss the biological sources of the three parts based on mathematical formulation.

It was proposed in ^8^ that collaboration between CpGs can be achieved by recruitment of methylase or demethylase enzymes, which akin to the situation of nucleosome modifications ^19^. Here, the power law decay suggests an alternative mechanism, by which the enzymes perform random walk along DNA and self-enhance dissociation. Accordingly, the enzyme concentration along the DNA can be described with a diffusive equation of form
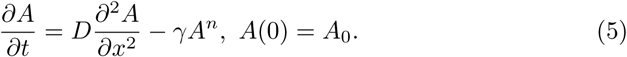

This equation gives the power-law decaying steady state solution of form *A* = *A*_0_(*x*/*ϵ* + 1)^−2/(*n*−1)^ in the case of self-enhance dissociation (*n* > 1) ^14^. Biologically, many experiments have revealed similar cohesion movement along DNA that play crucial roles in gene expression ^4,11^. Nevertheless, evidences for the molecular based of the correlation, either recruitment or movement along DNA, are waiting to further studies.

Interestingly, the bell-shape bulb is central at 160bp, right the distance of about one nucleosome. The strength coefficient *α* = 12 is consistent for early mouse embryo cells from sperm/oocyte to PGC E13.5 (Table 2), and the correlation is insensitive with changes in the parameter *α* (Figure 5). Moreover, various mouse and human somatic tissue cells give two catalogues cells with either small or large *α* values (Appendix C, Table C1). These observations suggest that the term 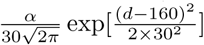 may represent a collaborative effect between DNA methylation and nucleosome modification, the collaboration strength *α* can be different for different type cells, but is insensitive with the kinetic parameters. A type of nucleosome modification, histone H3 lysine 9 dimethylation (H3K9me2), often show nucleation sites overlap to CGIs^2^, and is involved in DNA methylation^13,15,7^. Moreover, loss of DNA methylation enhances the removal of H3K3me3 under transcriptional stimulus^9^. These evidences support the correlation between nucleosome modification and DNA methylation, however the molecular mechanism is not yet clear.

Data analysis and model simulation have shown that the constant tail coefficient *c* is related to the global methylation level. Biologically, the global methylation level can be regulated by the enzyme activities that regulate DNA methylation. This observation remains further confirmation and theoretical consideration from the perspective of competition and collaboration of enzymes in methylation kinetics.

## 4. Conclusion

Inheritance of DNA methylation pattern is crucial during development. Here, driven from experimental data, we establish the formulation of the collaboration of DNA methylation. Applying the obtained function to a simple stochastic dynamic model can well reproduce the experimental observed correlation. The function reveal that there are three possible sources of the correlation: movement of enzymes along DNA, collaboration between DNA methylation and nucleosome modification, and global enzyme concentrations in the cell. Moreover, the collaboration strength between DNA methylation and nucleosome modification is universal in our study of mouse early embryo. These findings provide insightful understanding of the collaborations in DNA methylation, and for the mechanisms of inheritance of DNA methylation patterns. Nevertheless, despite the well support from experimental data, molecular details of the correlation in DNA methylation proposed here are waiting for further experimental studies.

## Acknowledgments

This work was supported by the National Natural Science Foundation of China (91430101 and 11272169).

## Appendix A. Correlation patterns are similar for different chromosomes in a cell

We calculated the correlation patterns of DNA methylation from the mouse embryo MethylC-Seq data ^17^. We choose the experimental data from the parental strain of mouse embryo at the 2-cell stage, and calculated the Pearson correlation of methylation levels between two adjacent CpG sites from chromosome 1 to 18. The relationship between the correlation and CpG distance are shown at Figure 1. From Figure 1, we can see that the correlation patterns are similar for different chromosomes.

**Fig. A1.**
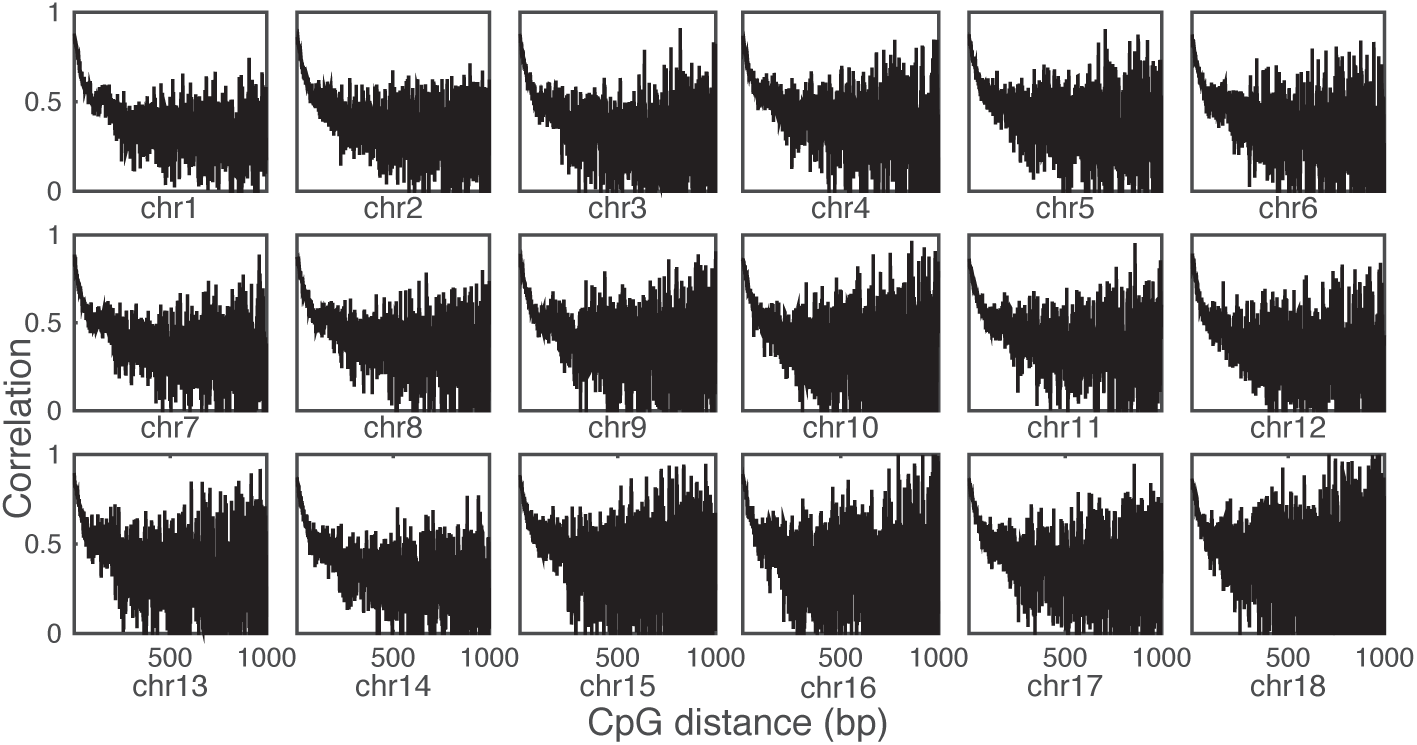
Pearson correlations, from chromosome 1 to 18, between DNA methylation levels of adjacent CpG sites as function of the distance (in bp) between CpG sites. Data obtained from the parental strain of mouse embryo at the 2-cell stage (P-2-cell-mouse, GEO: GSM1386021).

## Appendix B. Correlation patterns are different at different embryo stages

We calculated the correlation patterns of DNA methylation from different mouse embryo stages (Figure 2). The correlation patterns of paternal and maternal chromosomes are distinct to each other before E6.5, and tend to be consistent in latter stages.

## Appendix C. Correlation function for mouse and human tissues

Table C1 gives the coefficients and RMSD while we apply the procedure to experimental data of mouse and human tissues. Experimental data were obtained from the Whole Genome Bisulfite Sequencing (WGBS) data, GEO(ref. ^5^), and ENCODE(ref. ^3^). Fitting results are shown at Figure C1-C2.

**Fig. B1.**
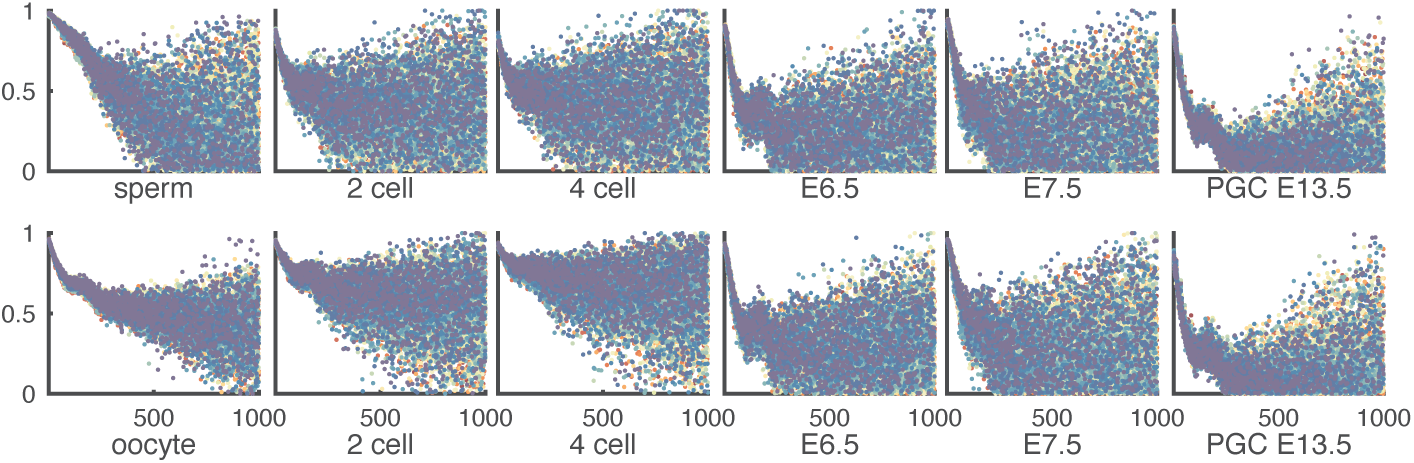
Pearson correlations between DNA methylation levels at adjacent CpG sites from different embryo stage of mouse (GEO: GSE56697). The correlation patterns of upper row are paternal chromosomes, and the bottom row are maternal chromosomes.

**Fig. C1.**
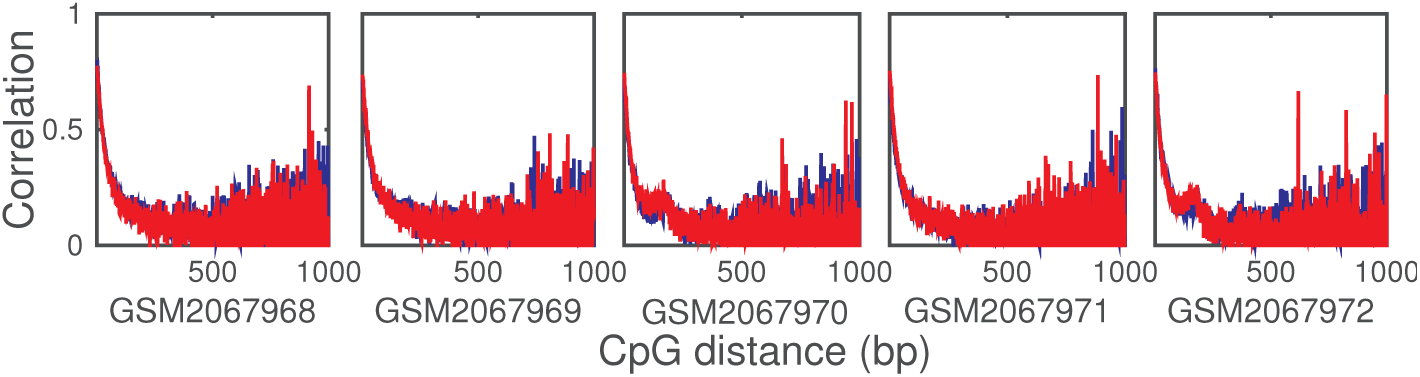
Correlations of mouse sample cells from Table C1. Blue for correlation obtained from experimental data, red for correlation obtained from models simulation.

**Fig. C2.**
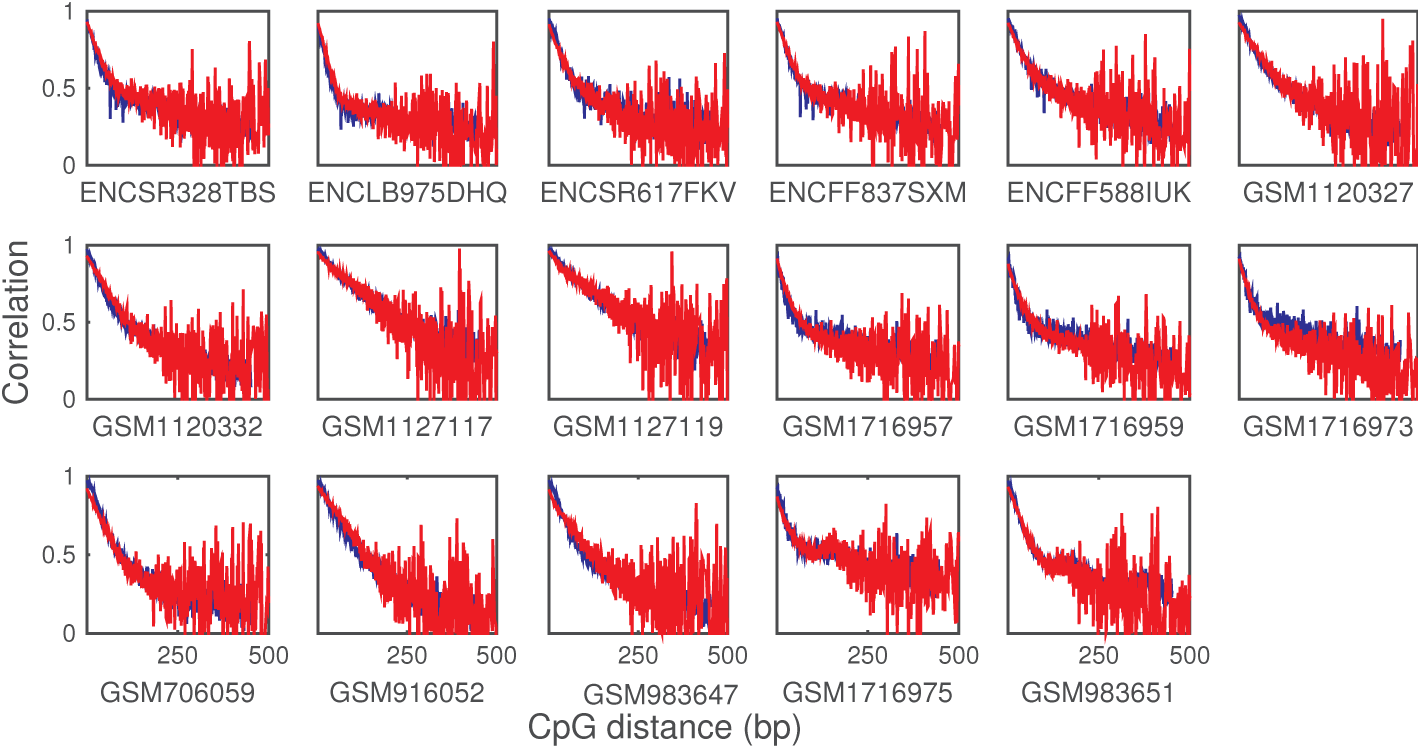
Correlations of human sample cells from Table C1. Blue for correlation obtained from experimental data, red for correlation obtained from models simulation.

**Table C1.**
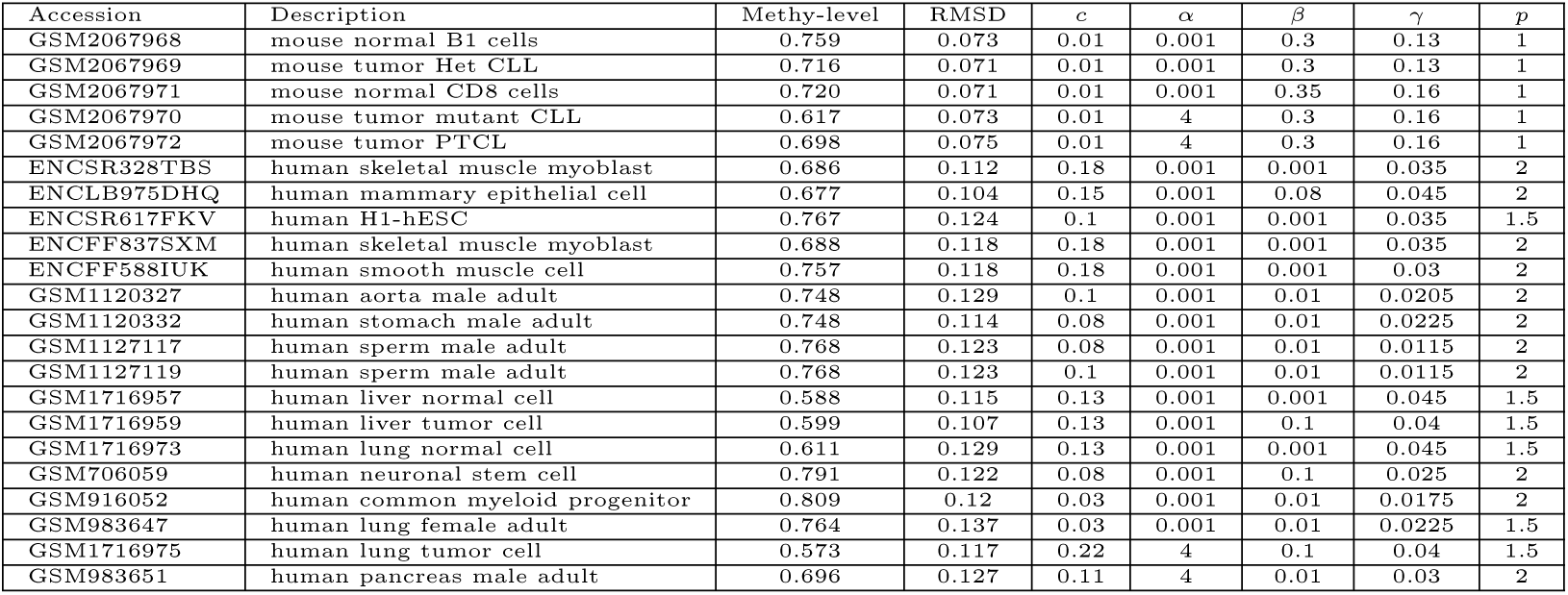
The data-source, average methylation levels and fitted coefficients of data.

